# Structural network alterations in focal and generalized epilepsy follow axes of epilepsy risk gene expression: An ENIGMA study

**DOI:** 10.1101/2021.10.18.464713

**Authors:** Sara Larivière, Jessica Royer, Raúl Rodríguez-Cruces, Maria Eugenia Caligiuri, Antonio Gambardella, Luis Concha, Simon S. Keller, Fernando Cendes, Clarissa L. Yasuda, Leonardo Bonilha, Ezequiel Gleichgerrcht, Niels K. Focke, Martin Domin, Felix von Podewills, Soenke Langner, Christian Rummel, Roland Wiest, Pascal Martin, Raviteja Kotikalapudi, Terence J. O’Brien, Benjamin Sinclair, Lucy Vivash, Patricia M. Desmond, Elaine Lui, Anna Elisabetta Vaudano, Stefano Meletti, Manuela Tondelli, Saud Alhusaini, Colin P. Doherty, Gianpiero L. Cavalleri, Norman Delanty, Reetta Kälviäinen, Graeme D. Jackson, Magdalena Kowalczyk, Mario Mascalchi, Mira Semmelroch, Rhys H. Thomas, Hamid Soltanian-Zadeh, Esmaeil Davoodi-Bojd, Junsong Zhang, Barbara A. K. Kreilkamp, Matteo Lenge, Renzo Guerrini, Khalid Hamandi, Sonya Foley, Theodor Rüber, Bernd Weber, Chantal Depondt, Julie Absil, Sarah J. A. Carr, Eugenio Abela, Mark P. Richardson, Orrin Devinsky, Mariasavina Severino, Pasquale Striano, Domenico Tortora, Sean N. Hatton, Sjoerd B. Vos, Lorenzo Caciagli, John S. Duncan, Christopher D. Whelan, Paul M. Thompson, Sanjay M. Sisodiya, Andrea Bernasconi, Angelo Labate, Carrie R. McDonald, Neda Bernasconi, Boris C. Bernhardt

## Abstract

Epilepsy is associated with genetic risk factors and cortico-subcortical network alterations, but associations between neurobiological mechanisms and macroscale connectomics remain unclear. This multisite ENIGMA-Epilepsy study examined whole-brain structural covariance networks in patients with epilepsy and related findings to postmortem co-expression patterns of epilepsy risk genes. Brain network analysis included 578 adults with temporal lobe epilepsy (TLE), 288 adults with idiopathic generalized epilepsy (IGE), and 1,328 healthy controls from 18 centres worldwide. Graph theoretical analysis of structural covariance networks revealed increased clustering and path length in orbitofrontal and temporal regions in TLE, suggesting a shift towards network regularization. Conversely, people with IGE showed decreased clustering and path length in fronto-temporo-parietal cortices, indicating a random network configuration. Syndrome-specific topological alterations reflected expression patterns of risk genes for hippocampal sclerosis in TLE and for generalized epilepsy in IGE. These imaging-genetic signatures could guide diagnosis, and ultimately, tailor therapeutic approaches to specific epilepsy syndromes.

## INTRODUCTION

Epilepsy is characterized by recurrent seizures and affects over 50 million people worldwide^1^. Cumulating evidence in epilepsy research has underscored the importance of interconnected brain networks in understanding the causes and consequences of the disease^2, 3^. In the common epilepsies, particularly temporal lobe epilepsy (TLE) and idiopathic generalized epilepsy (IGE), histopathological and neuroimaging studies have demonstrated structural and functional compromise across widespread brain networks^3, 4, 5, 6, 7^. Magnetic resonance imaging (MRI) analysis of brain morphology, including cortical thickness measurements and grey matter volumetry, provide *in vivo* evidence of structural alterations across multiple cortical and subcortical regions in both TLE^8, 9^ and IGE^10, 11^. Beyond small cohort studies, robust patterns of atrophy across widespread brain networks were identified in the common epilepsies through the ENIGMA-Epilepsy (<u>E</u>nhancing <u>N</u>euroImaging <u>G</u>enetics <u>t</u>hrough <u>M</u>eta-Analysis) consortium, with data aggregated from multiple international sites^12^.

Covariation of morphological MRI markers, termed “structural covariance analysis,” can extend earlier results on regional mapping of healthy and disease-related structural brain organization by identifying interregional network mechanisms. In healthy individuals, this approach has detected manifestations of functional crosstalk and structural connectivity between distributed regions^13, 14, 15^, and can tap into networks governed by shared genetic and maturational processes^16, 17, 18, 19, 20^. Prior work applying graph theoretical analyses to structural covariance networks has also characterized normative network topology^21^, revealing the presence of a “small world” organization. This architecture, which incorporates high clustering within segregated communities together with short paths between them, may provide a balance between network specialization and integration^22^. In TLE and IGE, structural covariance studies show syndrome-specific deviations from such a topological arrangement. In TLE, increased path length and clustering has been observed across both cortical systems^23, 24^ and in limbic/paralimbic^25^ networks. In contrast, diverging topological alterations have been reported in IGE, echoing either global increases in clustering^26, 27^ or path length^28^, global decreases in path length^27^, or no changes in network measures^29^. Analysis of structural brain metrics, gathered by ENIGMA-Epilepsy, provides an opportunity to consolidate network alterations in the common epilepsies in a generalizable manner.

Interactions across multiple spatial scales, ranging from genetic factors to macroscale cortical morphology and structural networks, shape cortical and subcortical organization in both health and disease^30^. When combined, these naturally intertwined dimensions offer new insights into the pathophysiology of system-level disorders such as epilepsy^31^. Neuroimaging studies of large-scale networks can profit from studies on the landscape of genetic risk factors in common epilepsies^32, 33^. Recently, the open release of *postmortem* human transcriptomics datasets, such as the Allen Human Brain Atlas (AHBA), has yielded opportunities to explore how gene co-expression patterns in the brain reflect macroscale neuroimaging findings^34, 35^. Integrating imaging and genetics can shed light on the micro-to-macroscale mechanisms that underly the pathophysiology of the common epilepsies. In parallel, this combination can also be used to understand the ways in which genes may drive, to some extent, network alterations in epilepsy. How, and whether, structural covariance network properties converge with spatial expression patterns of risk genes for epilepsies, however, remains an unanswered question.

In this ENIGMA-Epilepsy study, we aimed to identify robust structural network disruptions in individuals with TLE and IGE relative to healthy controls, aggregating inter-regional cortical thickness and subcortical volume correlations across 18 international sites. Graph theoretical analysis assessed global and regional topological disruptions in both epilepsy syndromes. Moreover, we leveraged gene expression information from the AHBA to relate macroscale network findings to spatial expression patterns of genetic risk factors in these two major forms of epilepsy. Spatial associations between topological changes and disease-specific gene expression maps were evaluated against null models with similar spatial autocorrelations^34, 36^. Reproducibility of our findings was also assessed across sites, variable network construction approaches, and clinical variables (side of seizure onset, disease duration).

## RESULTS

### Data samples

We studied 866 adult epilepsy patients (377 males, mean age±SD=33.82±9.48 years) and 1,328 healthy controls (588 males, mean age±SD=30.74±8.30 years) from 18 centres in the international Epilepsy Working Group of ENIGMA^37^. Our analyses focused on two patient subcohorts with site-matched healthy controls: TLE with neuroradiological evidence of hippocampal sclerosis (*n*_HC/TLE_=1,083/578, 257 right-sided focus) and IGE (*n*_HC/IGE_=911/288). Subject inclusion criteria and casecontrol subcohorts are detailed in the **Materials and Methods** and **Table 1**. Site-specific demographic and clinical information are provided in **Table S1**. All participants were aged between 18-50 years.

**TABLE 1.**
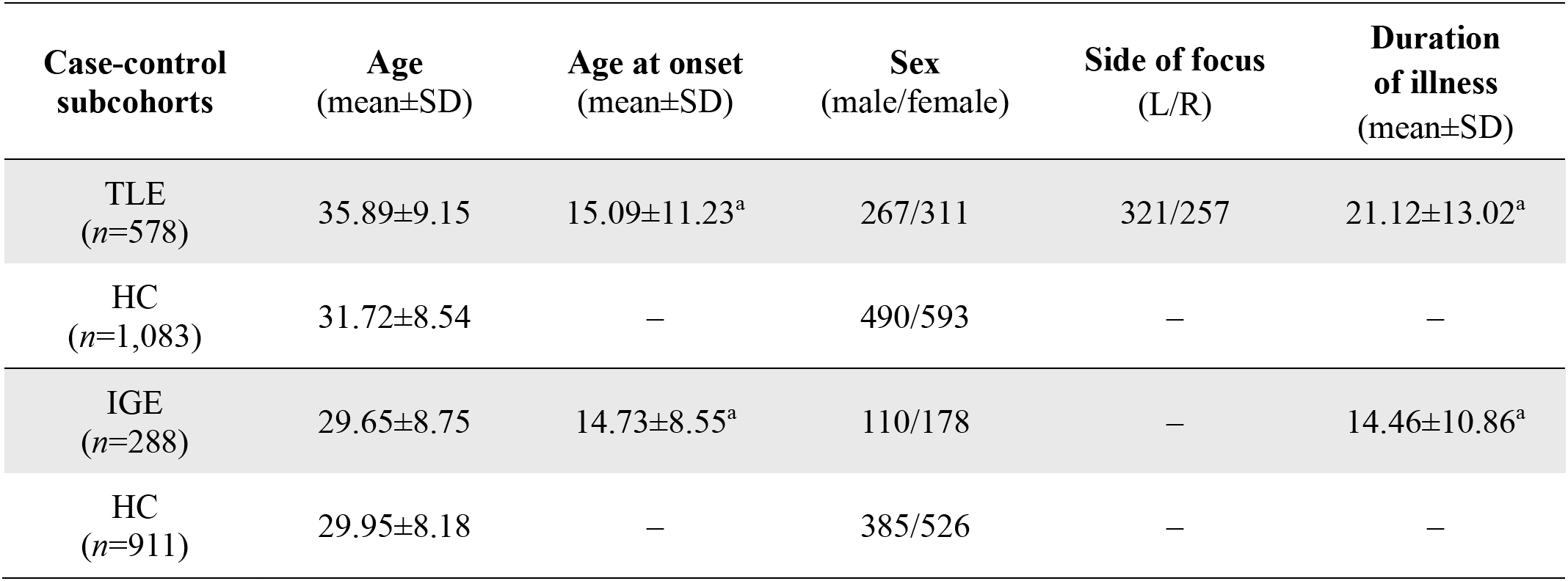
ENIGMA Epilepsy Working Group demographics. Demographic breakdown of patient-specific subcohorts with site-matched controls, including age (in years), age at onset of epilepsy (in years), sex, side of seizure focus (TLE patients only), and mean duration of illness (in years). Healthy controls from sites that did not have TLE (or IGE) patients were excluded from analyses comparing TLE (or IGE) to controls.^a^Information available in 544/578 TLE patients and 248/288 IGE patients.

### Global network characteristics

Cortical thickness (measured across 68 gray matter brain regions^38^) and volumetric data (measured across 12 subcortical gray matter regions and bilateral hippocampi) were obtained from all patients and controls. Cortical and subcortical data were harmonized across scanners and sites using CovBat^39^, and statistically corrected for age, sex, mean cortical thickness, and intracranial volume^39^. Group- and sitespecific structural covariance networks were then computed from morphometric (cortical thickness/subcortical volume) correlations (**Fig. 1A**). To characterize the topology of structural covariance networks, we computed two fundamental and widely used graph-theoretical parameters^40^: (*i*) mean clustering coefficient, to quantify local network efficiency, and (*ii*) mean path length, to index global efficiency (**Fig. 1B**).

**FIGURE 1.**
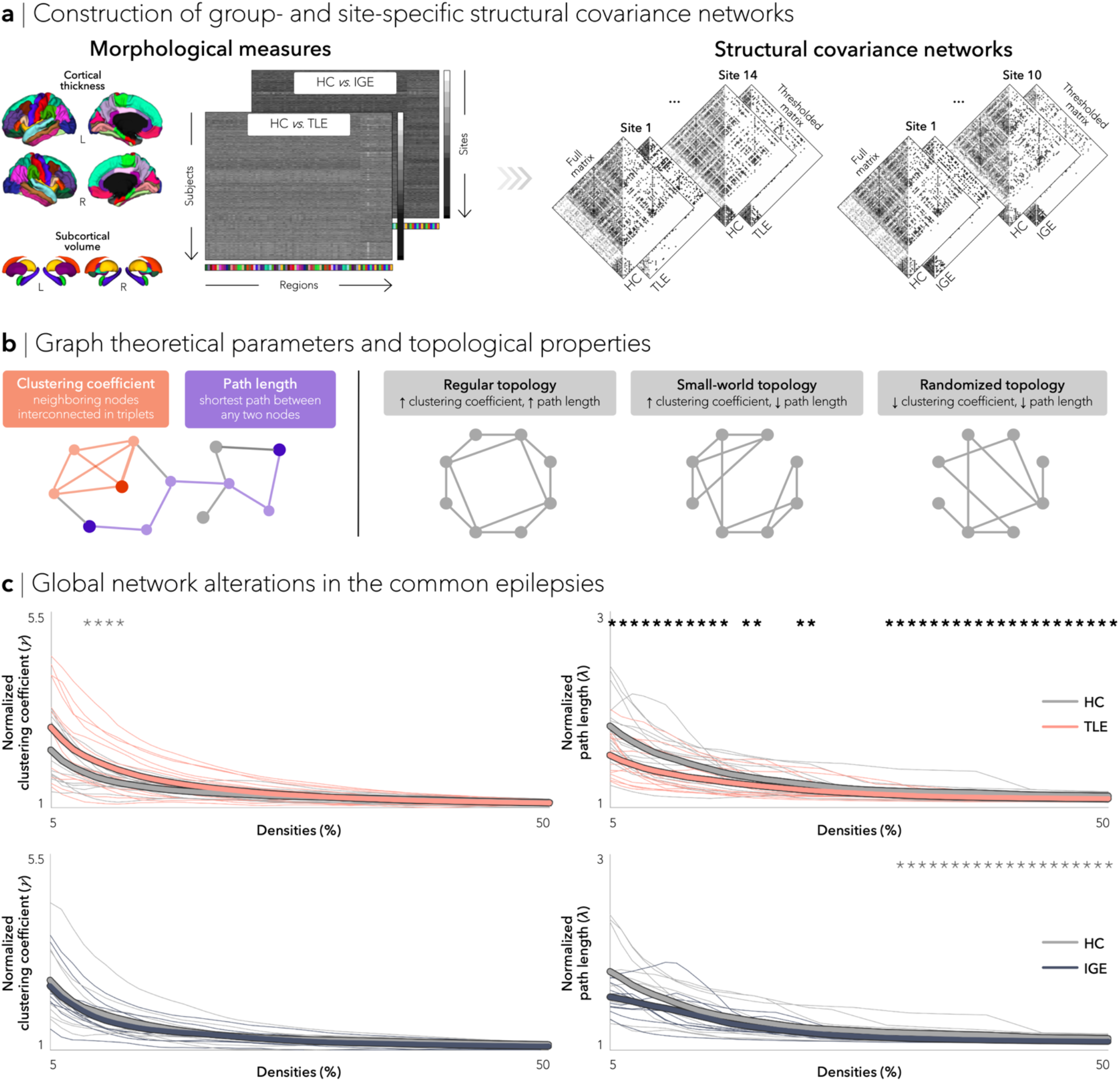
Structural covariance networks in the common epilepsies. (**a**) Schematic showing the construction of group- and site-specific structural covariance networks from morphometric correlations. (**b**) Two graph theoretical parameters characterized network topology: clustering coefficient, which measures connection density among neighboring nodes (orange) and path length, which measures the number of shortest steps between any two given nodes (purple). The interplay between clustering coefficient and path length can describe three distinct topological organizations: regular networks with high clustering and path length (left), small-world networks with high clustering and low path length (middle), and random networks with low clustering and path length (right). (**c**) Global differences in clustering coefficient (left) and path length (right) between TLE and HC (top) and between IGE and HC (bottom) are plotted as a function of network density. Increased small-worldness (*i.e*., increased clustering and decreased path length) was observed in individuals with TLE, whereas individuals with IGE showed decreases in clustering and path length, suggesting a more random configuration. Student’s t-tests were performed at each density value; bold asterisks indicate *p*_FDR_<0.1, semitransparent asterisks indicate *p*_uncorr_<0.05. Thin lines represent data from individual sites.

Comparing patients with TLE to controls, our multisite analysis revealed modest increases in overall clustering coefficient (*p*_uncorr_<0.05 at network densities (*K*)=0.08-0.11; see **Methods**) and decreases in mean path length over multiple density thresholds (*p*_FDR_<0. 1 at *K*=0.05–0.15, 17–18, 22–23, and 0.30–0.50) in patients. In contrast, IGE patients showed, on average, similar overall clustering coefficient relative to controls, but marginal decreases in overall path length (*p*_uncorr_<0.05 at *K*=0.31–0.50; **Fig. 1C**).

### Regional network characteristics

We also quantified clustering coefficient and path length changes at the nodal level. Multivariate comparisons, combining clustering coefficient and path length in TLE relative to controls, revealed trends for topological alterations in bilateral parahippocampus, paracentral lobule, lateral occipital cortex, putamen, and caudate, ipsilateral angular gyrus and orbitofrontal cortex, as well as contralateral insula, middle, and inferior temporal gyri (all *p*_FDR_<0.06). Although regional Cohen’s *d* effect sizes estimated across sites revealed an overall increase in small-worldness in TLE (Cohen’s *d* mean±SD: clustering=0.26±0.16, path length=-0.19±0.13), there were variations in different subnetworks. Notably, paralimbic and limbic regions such as the bilateral orbitofrontal, temporal, and angular cortices as well as ipsilateral amygdala showed an increase in clustering coefficient and path length (Cohen’s *d* mean±SD: clustering=0.20±0.13, path length=0.18±0.11), suggestive of a more regularized, “lattice-like,” subnetwork arrangement (**Fig. 2A**).

**FIGURE 2.**
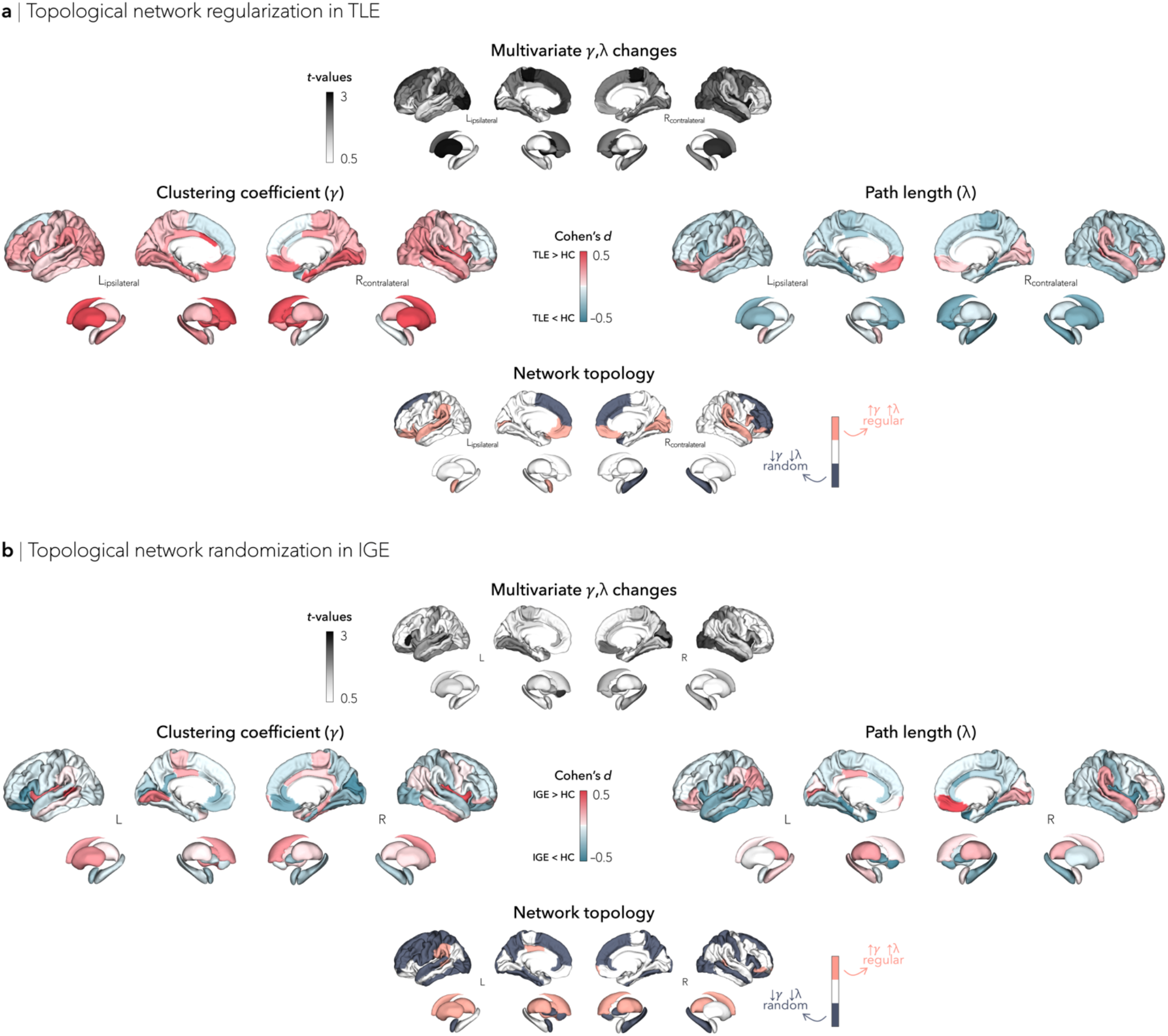
Nodal network alterations. (**a**) Graph theoretical analysis of structural covariance between individuals with TLE and controls revealed increased clustering and path length in bilateral orbitofrontal, temporal, and angular cortices, caudate, and putamen, as well as ipsilateral amygdala, revealing a regularized, “lattice-like,” arrangement. (**b**) In IGE, widespread multivariate topological alterations were observed in bilateral fronto-temporo-parietal cortices, right nucleus accumbens, and left pallidum. Clustering and path length effect sizes in these regions suggest a randomized network configuration (decreased clustering and path length).

When compared to controls, individuals with IGE showed widespread multivariate topological alterations in left inferior frontal gyrus *pars opercularis*, superior temporal sulcus, and nucleus accumbens, and right calcarine sulcus, insula, inferior temporal gyrus, and lateral occipital cortex (all *p*_FDR_<0.013). Effect sizes for each individual metric revealed decreased clustering coefficient and path length, with predominant changes in bilateral fronto-temporo-parietal cortices, nucleus accumbens, and pallidum (Cohen’s *d* mean±SD: clustering=-0.15±0.11, path length=-0.22±0.17), suggesting a more randomized network configuration (**Fig. 2B**). Increased small-worldness in IGE was also observed in fronto-parietal (bilateral paracentral lobule, right precentral gyrus) and temporal (left middle temporal gyrus, right inferior temporal gyrus) regions, but appeared to be primarily driven by conspicuously shorter path length (Cohen’s *d* mean±SD: clustering=0.16±0.14, path length=-0.27±0.19).

### Transcriptomic associations

Having established multivariate topological abnormalities in TLE and IGE, we evaluated whether these network-level findings were associated with the spatial expression patterns of previously established genetic risk factors. To this end, we assessed spatial correlations between epilepsy-related gene expression maps (genes listed in a recently published GWAS^32^ and co-expression levels obtained from the Allen Human Brain Atlas^34^; see **Materials and Methods** and **Table S2**) and multivariate topological profiles. To assess specificity, we also cross-referenced our network findings with five additional transcriptomic maps from common neuropsychiatric conditions, including: attention deficit/hyperactivity disorder^41^, autism spectrum disorder^42^, bipolar disorder^43^, major depressive disorder^44^, and schizophrenia^45^. Significance of spatial correlations was established using spin permutation tests^36^ that control for spatial autocorrelations (see **Materials and Methods**), from the ENIGMA Toolbox [https://github.com/MICA-MNI/ENIGMA;^46^].

We found significant associations between the spatial patterns of multivariate topological alterations in TLE and epilepsy risk gene co-expression levels of hippocampal sclerosis (*r*=0.21, *p*_spin_<0.05). On the other hand, multivariate topological differences in IGE were preferentially related to the co-expression levels of generalized epilepsy (*r*=0.20, *p*_spin_<0.05; **Fig. 3A**). In both TLE and IGE, network-level findings did not correlate with other disease-related transcriptomic maps (range *r*=-0.047–0.12, all *p*_spin_>0.1; **Fig. 3B**).

**FIGURE 3.**
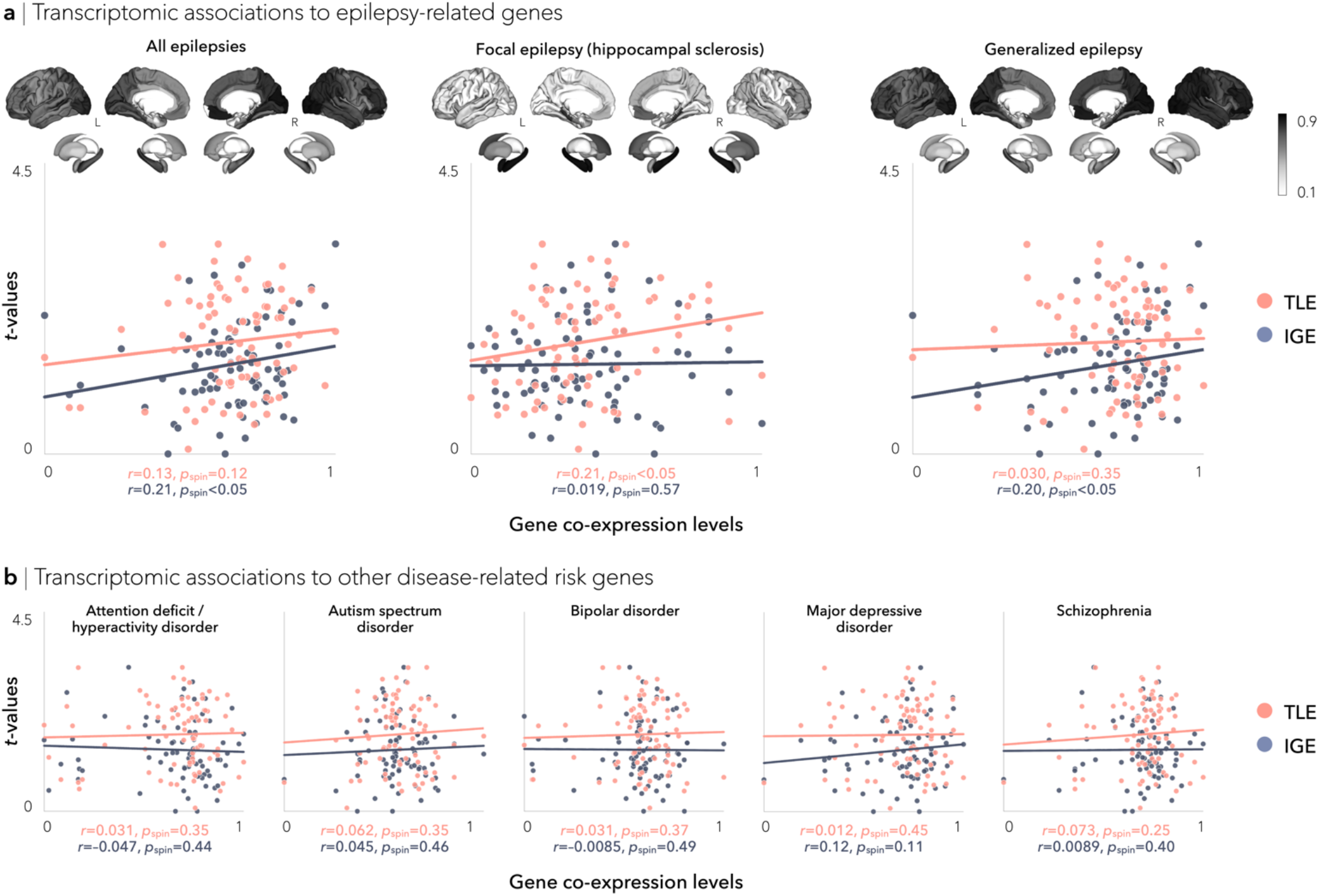
Relations between epilepsy gene co-expression and network topology. (**a**) Epilepsy-related gene coexpression levels are mapped to cortical and subcortical surface templates. Spatial associations between microarray data and multivariate topological changes were strongest for expression levels of hippocampal sclerosis genes in TLE (*r*=0.21, *p*_spin_<0.05) and generalized epilepsy genes in IGE (*r*=0.20, *p*_spin_<0.05). (**b**) Network associations did not correlate with other disease-related transcriptomic maps, suggesting specificity of these associations to epilepsy-related molecular risk factors.

### Associations to clinical variables

As seizure focus laterality may differentially affect structural covariance networks^23^, we repeated the above analyses in left and right TLE independently, comparing patient subgroups to both controls and to each other. Global increases in clustering and decreases in path length were observed in both left (clustering: *p*_FDR_<0.05 at *K*=0.05–0.26. 28–29; path length: *p*_FDR_<0.05 at *K*=0.05. 0.07–0.22. 25–39; **Fig. S1A**) and right (clustering: *p*_FDR_<0.05 at *K*=0.05–0.25; path length: *p*_uncorr_<0.05 at *K*=0.14–0.17; **Fig. S2A**) TLE patients relative to controls. Similarly, dominant patterns of multivariate (clustering coefficient and path length) topological changes in bilateral lateral occipital cortex (*p*_FDR_<0.01), parahippocampus (*p*_FDR_<0.005), entorhinal cortex (*p*_FDR_<0.01), and insula (*p*_uncorr_<0.05), ipsilateral precuneus (*p*_FDR_<0.01), anterior cingulate cortex (*p*_FDR_<0.01), and superior temporal gyrus (*p*_FDR_<0.05), as well as contralateral middle temporal gyrus (*p*_FDR_<0.05) were observed when comparing left (**Fig. S1A**) and right (**Fig. S2A**) TLE cohorts separately to controls. Left TLE additionally showed alterations in bilateral paracentral and sensorimotor cortices (*p*_FDR_<0.01), while right TLE additionally showed abnormalities in bilateral hippocampi (*p*_FDR_<0.005). Direct comparison of left *vs*, right TLE revealed no significant global (*p*_FDR_>0.27; **Fig. S3A**) nor regional (*p*_FDR_>0.11; **Fig. S3B**) differences between the two subcohorts. Effect sizes for clustering and path length indicated network regularization of the mesiotemporal and postcentral gyrus subnetwork in left TLE (**Fig. S1A**), but widespread cortical regularization in right TLE (**Fig. S2A**); these slight differences in regional topological configurations were further confirmed in left *vs*, right TLE comparisons (**Fig. S3B**). Differences in multivariate topological changes between left and right TLE marginally affected their associations to disease-related risk genes; in both groups, strongest spatial correlations were observed with co-expression levels of genes previously associated to hippocampal sclerosis (**Fig. S1B. Fig. S2B**).

We repeated the structural covariance analyses in patients grouped by duration of illness using a median split approach (TLE=20 years, *n*_short-TLE_=270. *n*_long-TLE_=275; IGE=15 years, *n*_short-IGE_=137. *n*_long-IGE_=111). In TLE and IGE, both patient subgroups (short and long duration) showed similar patterns to the overall between-group differences when compared to controls (TLE: **Fig. S4A**; IGE: **Fig. S5A**). Nevertheless, in TLE, we observed a shift in network regularization from fronto-central and limbic regions (shorter duration) to frontotemporal and limbic regions (longer duration; **Fig. S4B**). Conversely, in IGE, we observed both network randomization (fronto-central regions) and regularization (fronto-parietal regions) in patients with short and long duration. Direct comparison of patients with short *vs*, long duration of TLE or IGE revealed no significant global (TLE: *p*_FDR_>0.5. **Fig. S6A**; IGE: *p*_FDR_>0.20. **Fig. S6B**) nor regional (TLE: *p*_FDR_>0.072. **Fig. S6A**) differences between pairs of subcohorts, with the exception of patients with shorter duration of IGE showing multivariate topological changes in bilateral fronto-limbic areas relative to those with longer duration (*p*_FDR_<0.05. **Fig. S6B**).

### Robustness of findings across different sites and analysis thresholds

Despite some site-to-site variability, syndrome-specific global structural covariance differences were overall consistent across sites and similar to those obtained from the multisite aggregation for both TLE and IGE patients (**Fig. 1C**). As observed in the multisite findings, sitespecific increases in clustering coefficient and path length in TLE were most frequently observed in orbitofrontal, temporal, and angular cortices as well as amygdala (**Fig. S7A**). Similarly, in agreement with the multisite findings, site-specific decreases in clustering and path length in IGE were most consistent in frontoparietal cortices and hippocampus (**Fig. S7B**).

Our findings were not affected by varying the density of structural covariance networks: Across the range of possible thresholds, we observed high correlations among multivariate topological brain maps computed from thresholded structural covariance matrices in TLE (95.35% of correlations were below *p*_spin_<0.1) and IGE (90.86% of correlations were below *p*_spin_<0.1; **Fig. S8A**). Moreover, we observed comparable associations between topological abnormalities (computed across the range of thresholds) and gene co-expression levels, with highest stability in TLE observed for correlations of topological alterations and focal epilepsy with hippocampal sclerosis gene co-expression levels (62.00% of correlations were below *p*_spin_<0.1). Conversely, stability in IGE was highest for correlations with gene co-expression levels of generalized epilepsy (40.00% of correlations were below *p*_spin_<0.1), major depressive disorder (42.00% of correlations were below *p*_spin_<0.1), and schizophrenia (52.00% of correlations were below *p*_spin_<0.1; **Fig. S8B**).

## DISCUSSION

This worldwide ENIGMA study is the largest investigation of structural covariance networks in the common epilepsies and bears robust evidence for syndrome-specific topological disruptions. First, despite showing global increases in small-worldness in TLE as compared to controls, profound regional alterations in orbito-fronto-temporal regions indicated a shift towards a more regularized, “lattice-like,” subnetwork configuration. In contrast, IGE presented with widespread decreases in clustering and path length in fronto-temporo-parietal cortices, indicating a more random topology. These syndrome-specific networklevel findings were spatially related to the expression pattern of genetic risk factors associated with hippocampal sclerosis and generalized epilepsy in recent genome wide association studies (GWAS)^32^. Findings were highly consistent across sites and methodologies, corroborating robustness and generalizability. Taken together, our study identifies imaging-genetic signatures in the common epilepsies, which ultimately, may facilitate early diagnosis and lead to the development of new and improved treatment strategies.

We performed graph theoretical analysis on MRI-based cortical thickness and subcortical volume correlations in adults with TLE, adults with IGE, and healthy controls^19, 47^. Our covariance analysis extends prior research on atrophy mapping by tapping into the topology of inter-regional structural brain networks and describing the network organization underlying wholebrain pathological interactions in the common epilepsies. Using a multisite approach, we showed that patients with TLE preserved an overall small-world configuration with increased clustering and decreased path length over a wide sparsity range. Upon examination of uni- and multivariate regional changes, however, we found key differences between distinct brain subnetworks. Topological alterations were most marked in a subnetwork comprising orbitofrontal, temporal, and angular cortices, pointing to increased local connectivity (*i.e*., a more regular configuration) in TLE than in healthy controls. This bilateral topological regularization was observed in both left and right TLE patients, albeit more constrained to fronto-temporal cortices in left TLE, a difference that may be attributable to asymmetrical structural damage or to higher connectivity of the dominant hemisphere^48^. Notably, when split into short *vs*, long duration groups, patients with longer duration of TLE exhibited topological regularization primarily in temporo-parietal cortices, areas that show progressive pathological changes unrelated to normal aging^3, 49, 50, 51^. Although group-level alterations in the hippocampus were modest, with right TLE patients displaying slightly more severe abnormalities than left TLE, intrinsic hippocampal deafferentation may nevertheless contribute to extrahippocampal reconfigurations, affecting neighbouring regions including orbitofrontal and temporal cortices, as well as the amygdala^52^. Given the high density of connections from the hippocampus to the rest of the brain^53, 54^, neuronal loss and deafferentation within limbic structures may cause local excess connectivity and decreased internetwork covariance in remote regions. Such a topological shift may be supported by findings in animal models^55^ as well as human diffusion MRI^52, 56, 57^ and functional connectivity distance studies^58^, which have highlighted imbalances in short- *vs*, long-range connections in epilepsy-related pathology. More regularized networks are spatially compact, which may facilitate recurrent excitatory activity and high frequency oscillations, and may be attributable to a loss of temporo-limbic structural connections^55, 59^. In line with prior EEG/intracranial EEG studies, network regularization has often been reported at seizure onset, a configuration that shifts toward a globally integrated process as the seizure spreads, eventually reaching a random configuration upon seizure termination^60^. Understanding such structural reorganization offers a comprehensive knowledge of the neural substrates and pathophysiological mechanisms of TLE.

In contrast with TLE, overall structural covariance network configurations in IGE showed a tendency away from aberrant local connectivity and towards a more random architecture. On the whole, increased randomness of the brain’s structural network organization denotes reduced local efficiency but increased global efficiency^21, 61^. These global topological findings were complemented by regionspecific mapping of graph-theoretical parameters, which also identified widespread regional alterations in IGE patients. Although the pattern was overall mixed across regions—*a fortiori* in smaller patient subgroups split according to disease duration—showing increases and decreases in both path length and clustering, a large subnetwork comprising frontal, temporal and parietal cortices showed concomitant reductions in path length as well as clustering. A decrease in clustering implies reductions in local specialization, but a decrease in path length (*i.e*., increased global network efficiency) may indicate an imbalance in the integration and segregation of structural covariance network organization. Such imbalance might explain, at least in part, the ability of seizures to rapidly spread, not just locally but in a diffuse manner within bilateral fronto-temporo-parietal cortices in IGE patients^62^. In rodent models of IGE, frontoparietal cortices have typically shown increased simultaneous neuronal activity during generalized seizures^63, 64^ Moreover, extensive neuroimaging work in IGE patients that analyzed cortical morphology has shown widespread cortical structural network compromise^6, 65^, with midline frontal and paracentral regions emerging as potential epicenters of morphological abnormalities in IGE^3, 66^. Notably, IGE patients also presented with focal patches of network randomization, similar to TLE patients in paralimbic subnetworks. Affected regions included paralimbic cortices, but coupled with subcortical structures, notably the thalamus. Extensive evidence supports atypical thalamo-cortical interactions as being at the core of the pathophysiological network of IGE^7, 10, 11, 67, 68^, with aberrant thalamo-cortical loops contributing to the generation of spike and slow wave discharges^69^. Alterations of thalamic morphology and metabolism, as well as of its functional and structural connectivity with widespread cortical networks, have also been reported in a convergent neuroimaging literature across several IGE syndromes^10, 70, 71^. In future work, it will be of interest to explore the consistency, or variability, of these topological imbalances in thalamic as well as cortical subnetworks are across different IGE subsyndromes. It is also important to understand how they vary with respect to other clinically relevant parameters including levels of drug-control. In that context, we recommend further increasing the spatial resolution, allowing for a fine-grained assessment of both cortical network architecture and thalamic subdivisions. This could be achieved, for example, by adopting recent approaches that reported structural, functional, and microcircuit anomalies in IGE compared to both TLE patients and healthy controls^6, 7^.

Connectome topology has been extensively studied in healthy and diseased brains, however, research investigating associations between macroscale findings and the genetic architecture of epilepsy is still in infancy. A recent genome-wide mega-analysis performed in the common epilepsies identified 21 biological candidate genes across 16 risk loci, thus providing initial evidence for epilepsy-associated gene expression changes^32^. By integrating neuroimaging and transcriptional atlas data, here we tested the hypothesis that transcriptomic vulnerability would covary with structural network abnormalities in TLE and IGE. We showed that epilepsy-related variations in brain network topology spatially converged with gene expression profiles of risk genes for each syndrome. Specificity of these associations was further evidenced by the fact that topological alterations did not correlate with transcriptional signatures of several common psychiatric disorders. In the long term, these imaging-genetic associations may form the foundation for translation and clinical studies aiming to tailor therapeutic approaches to specific epilepsy syndromes. In parallel, our results may serve in the development of more effective treatments that can be targeted to the individual patients based on their genetic profile. Some limitations of the gene expression associations must be highlighted, including the fact that our understanding of the genetic architecture of the epilepsies is evolving, with the reported risk genes likely being expanded and refined as more genetic data in epileptic patients become available. Moreover, the gene expression information in the current work was derived from a small sample of six *postmortem* human donor brains, with predominant cortical and subcortical genetic microarray sampling performed in the left hemisphere. Even so, our findings suggest that genes previously associated with specific epilepsy syndromes were over-represented in regions that share similar topological alterations. In keeping with prior molecular studies in epilepsy^72, 73, 74, 75^, we speculate that differentially expressed epilepsy-related gene sets may contribute to a selective vulnerability of networks for structural reconfigurations in TLE and IGE. Replication of these distinctive imaging-genetic signatures in more comprehensive gene expression datasets may thus hold significant promise for stratification and effective treatment of epilepsy. Exploiting individualized gene expression profiles in the same cohort of patients, therefore, seems to be the logical next step to improve imaging-genetic associations and update our understanding of causes and consequences of epilepsy.

Several sensitivity analyses suggested that our findings were not affected by differences in scanners or sites or methodological choices. Site and scanner effects were mitigated for the most part using CovBat, a postacquisition statistical batch normalization process used to harmonize between-site and between-protocol effects in mean, variance, and covariance, while protecting biological covariates (*e.g*., disease status)^39^. Multivariate topological findings as well as associations between network-level findings and gene expression maps were consistently observed across different matrix thresholds. Despite some site-to-site variability in global and regional graph theoretical metrics, findings were overall similar across independent centers and reflected those from the multisite aggregation. As data sharing practices can at times be challenging, in part due to privacy and regulatory protection. ENIGMA represents a practical alternative for standardized data processing and anonymized derivative data^12, 76, 77, 78, 79, 80^. This collaborative effort allowed us to identify a robust association of brain structural network changes with patterns of expression of genetic risk factors in the common epilepsies, while addressing robustness of effects across clinical subgroups, international sites, and methodological variations. The imaging-genetic associations identified herein could guide diagnosis of common epilepsies, and ultimately, contribute to the development of tailored, individualized, and syndrome-specific therapeutic approaches.

## MATERIALS AND METHODS

### ENIGMA participants

Epilepsy specialists at each center diagnosed patients according to the seizure and syndrome classifications of the International League Against Epilepsy (ILAE). Inclusion of adults with TLE was based on the combination of electroclinical features and MRI findings typically associated with underlying hippocampal sclerosis. Inclusion of adults with IGE was based on the presence of tonic-clonic, absence, or myoclonic seizures with generalized spike-wave discharges on EEG. We excluded participants with a progressive or neurodegenerative disease (*e.g*., Rasmussen’s encephalitis, progressive myoclonus epilepsy), malformations of cortical development, tumors, or prior neurosurgery. Local institutional review boards and ethics committees approved each included cohort study, and written informed consent was provided according to local requirements.

### Cortical thickness and subcortical volume data

All participants underwent structural T1-weighted brain MRI scans at each of the 18 participating centers, with scanner descriptions and acquisition protocols detailed elsewhere^37^. Images were independently processed by each center using the standard ENIGMA workflow. In brief, models of cortical and subcortical surface morphology were generated with FreeSurfer 5.3.0^81^. Based on the Desikan-Killiany anatomical atlas^38^, cortical thickness was measured across 68 grey matter brain regions and volumetric measures were obtained from 12 subcortical grey matter regions (bilateral amygdala, caudate, nucleus accumbens, pallidum, putamen, thalamus) as well as bilateral hippocampus. Missing cortical thickness and subcortical volume data were imputed with the mean value for that given region; participants with missing data in at least half of the cortical or subcortical brain measures were excluded.

Data were harmonized across scanners and sites using CovBat—a batch-effect correction tool that uses a Bayesian framework to improve the stability of the parameter estimates^39^. Cortical thickness and volumetric measures were corrected for age and sex. Residualized data were *z*-scored relative to site-matched pooled controls and sorted into measures that were ipsilateral/contralateral to the focus.

### Covariance networks

Covariance networks were computed from cortical thickness and subcortical volume correlations. Interregional association matrices were first generated for each group (TLE, HC_TLE_, IGE, HC_IGE_) and each site with at least 10 participants per diagnostic group (*n*_TLE/HC_=15 sites, *n*_GE/HC_=10 sites), resulting in a total of 50 covariance matrices (*R*). In each matrix *R*, an individual entry *Ri,j* (with regions *i* and *j*) contained the pairwise linear product-moment cross-correlation coefficient of structural morphometry across group- and site-specific subjects.

### Network thresholding

Prior to analysis, negative correlations were set to zero and covariance network matrices were thresholded (density range of *K*=0.05–0.50, density interval of 0.01). This approach ensured that networks in all groups had an identical number of edges^82^ and that group differences were not primarily driven by low-level correlations^83^.

Among the network density levels of *K*=0.05–0.50, the network connectedness criterion (≥75% of nodes remain connected to other nodes within the network in at least 90% of sites) was satisfied only in the narrower *K*=0.08–0.50 range. For the main regional topological analyses, structural networks were constructed at a density of *K*=0.08.

### Global network properties

From the thresholded structural covariance networks, we computed two global metrics^84^: (*i*) mean clustering coefficient, which quantifies the tendency for brain regions to be locally interconnected with neighboring regions, and (*ii*) mean path length, which quantifies the mean minimum number of edges (*i.e*., connection between two regions) that separate any two regions in the network. Each measure was normalized relative to corresponding measures from 1,000 randomly generated networks with similar degree and weight properties, and subsequently averaged across all cortical and subcortical regions, separately.

### Regional network properties

Regional differences in topological parameters were assessed using an approach similar to the global network analysis; from the thresholded structural covariance networks, normalized clustering coefficient and normalized path length metrics were computed for every cortical and subcortical brain region. Individual nodal network parameters in patients were compared to controls across sites via Cohen’s *d* effect sizes. From effect size maps, topological profiles were generated to reflect either network regularization (areas of increased clustering coefficient and path length) or randomization (areas of decreased clustering coefficient and path length)^85^. Multivariate surface-based linear models were subsequently used to compare the aggregate of clustering coefficient and path length differences in patients relative to controls.

### Transcriptomic associations

The Allen Institute for Brain Science released the AHBA—a brain-wide gene expression atlas comprising microarray expression data from over 20,000 genes sampled across 3,702 spatially distinct tissue samples collected from six human donor brains^34^. Microarray expression data were first generated using *abagen*^86^, a toolbox that provides reproducible workflows for processing and preparing gene co-expression data according to previously established recommendations^87^. Preprocessing steps included intensity-based filtering of microarray probes, selection of a representative probe for each gene across both hemispheres, matching of microarray samples to brain parcels from the Desikan-Killiany atlas^38^, normalization, and aggregation within parcels and across donors. Genes whose similarity across donors fell below a threshold (*r*<0.2) were removed, leaving a total of 12,668 genes for analysis.

Leveraging a recently published GWAS from the International League Against Epilepsy Consortium on Complex Epilepsies^32^, we extracted the most likely genes associated with significant genome-wide loci in the common epilepsies (*i.e*., all epilepsies, *n*_genes_=20, as well as in two epilepsy syndromes (*i.e*., focal epilepsy with hippocampal sclerosis, *n*_genes_=3 and generalized epilepsy, *n*_genes_=13). We also queried additional lists of disease-related genes (obtained from other recently published GWAS), including gene sets for attention deficit/hyperactivity disorder [*n*_genes_=17^41^], autism spectrum disorder [*n*_genes_=22^42^], bipolar disorder [*n*_genes_=27^43^], major depressive disorder [*n*_genes_=194^44^], and schizophrenia [*n*_genes_=150^45^]. All gene sets were mapped to cortical and subcortical regions using the Allen Human Brain Atlas^34^ and projected to surface templates.

### Spatial permutation tests

The intrinsic spatial smoothness in two given brain maps may inflate the significance of their spatial correlation, if the spatial dependencies in the data are not taken into account^36^. Statistical significance of spatial correlations (*e.g*., between multivariate topological patterns and transcriptomic maps) was assessed using spin permutation tests^46^. This framework generates null models of overlap between cortical maps by projecting the spatial coordinates of cortical and subcortical data onto the surface spheres (*i.e*., the parameter spaces), applying randomly sampled rotations (10,000 repetitions unless specified otherwise), and reassigning original values^36^. The empirical correlation coefficients are then compared against the null distribution determined by the ensemble of spatially permuted correlation coefficients.

### Associations to clinical variables

As seizure focus lateralization may differentially impact topological organization of structural covariance networks^24^, we repeated the graph theoretical and transcriptomics analyses by comparing (*i*) left (*n*_LTLE_=321) and right (*n*_RTLE_=257) TLE independently to controls and (*ii*) directly comparing left *vs*, right TLE.

To study the effects of duration of illness on structural covariance networks, we repeated the graph theoretical analyses by comparing (*i*) patients with short duration of TLE or IGE (TLE duration<20 years, *n*_short-TLE_=270; IGE duration<15 years, *n*_short-IGE_=137) and patients with long duration of TLE or IGE (TLE duration≥20 years, *n*_long-TLE_=275; IGE duration≥15 years, *n*_long-IGE_=111) independently to controls, (*ii*) directly comparing short *vs*, long TLE, and (*iii*) directly comparing short *vs*, long IGE. Median splits were used to group short *vs*, long duration patients.

### Reproducibility and sensitivity analyses

#### Reproducibility across sites

To address reproducibility of our findings across different sites, we repeated our multivariate topological analysis independently in each site (*n*_TLE/HC_=14 sites, *n*_IGE/HC_=10 sites).

#### Stability across matrix thresholds

To verify that results were not biased by choosing a particular threshold, we repeated the network analyses and associations with disease-related gene co-expression levels across the range of matrix thresholds (*K*=0.05–0.50 with increments of 0.01). Specifically, Hotelling’s T^2^ multivariate (clustering coefficient and path length) topological changes comparing patients to controls were computed from structural covariance networks thresholded at every density and pairwise spatial correlations between all pairs of multivariate brain maps were performed. Spatial correlations between density-specific multivariate topological alterations and all disease-related transcriptomic maps were also assessed. Significance testing of these correlations was assessed via spin permutation tests with 1,000 repetitions.

## ACKNOWLEDGEMENTS

The authors would like to express their gratitude to the open science initiatives that made this work possible: (*i*) the ENIGMA Consortium (core funding for ENIGMA was provided by the NIH Big Data to Knowledge (BD2K) program under consortium grant U54 EB020403 to P.M.T.) and (*ii*) The Allen Human Brain Atlas and the abagen toolbox (https://doi.org/10.5281/zenodo.4984124).

Sara Larivière was funded by the Fonds de la Recherche du Québec – Santé (FRQ-S), Canadian Institutes of Health Research (CIHR), and the Ann and Richard Sievers Neurosciene Award, Raul Rodríguez-Cruces was funded by the FRQ-S, Jessica Royer was supported by a Canadian Open Neuroscience Platform (CONP) fellowship and CIHR, Lorenzo Caciagli acknowledges support from a Berkeley Fellowship jointly awarded by UCL and Gonville and Caius College, Cambridge, and by Brain Research UK (award 14181). The UNAM site was funded by UNAM-DGAPA (IB201712, IG200117) and Conacyt (181508 and Programa de Laboratorios Nacionales). Mark Richardson was funded by UK Medical Research Council grant MR/K013998/1. Fernando Cendes and Clarissa Yasuda were supported by the São Paulo Research Foundation (FAPESP). Grant # 2013/07559-3 (BRAINN - Brazilian Institute of Neuroscience and Neurotechnology). Stefano Meletti and Anna Elisabetta Vaudano were supported by the Ministry of Health (MOH). grant # NET-2013-02355313, Paul Thompson was funded by R01 NS106957 and P41 EB015922, Carrie R. McDonald was supported by ENIGMA-R21 (NIH/NINDS R21NS107739) and R01 (NIH/NINDS R01 NS122827), Sanjay M. Sisodiya was supported by the Epilepsy Society. UK. Boris C. Bernhardt acknowledges research support from the National Science and Engineering Research Council of Canada (NSERC Discovery-1304413), the CIHR (FDN-154298, PJT-174995). SickKids Foundation (NI17-039). Azrieli Center for Autism Research (ACAR-TACC), BrainCanada, FRQ-S, and the Tier-2 Canada Research Chairs program.

## ETHICS DECLARATIONS

Paul M. Thompson is funded in part by a research grant from Biogen, Inc. for work unrelated to the current manuscript.

## Supplementary Information

**TABLE S1.** Site-specific group demographics. Site-specific demographic breakdown of patient-specific subcohorts, including age (in years), age at onset of epilepsy (in years), sex, side of seizure focus (TLE patients only), and mean duration of illness (in years). TLE=Temporal lobe epilepsy, IGE=idiopathic/genetic generalized epilepsy, HC=healthy controls.

| Site name | Case-controls subcohorts | Age (mean±SD) | Age at onset (mean±SD) | Sex (male/female) | Side of focus (L/R) | Duration of illness (mean±SD) |
| --- | --- | --- | --- | --- | --- | --- |
| BERN | TLE *n*=18 | 31.28±9.09 | — | 9/9 | 10/8 | — |
|  | IGE *n*=11 | 31.36±8.79 | — | 5/6 | — | — |
|  | HC *n*=72 | 31.43±8.44 | — | 34/38 | — | — |
| BONN | TLE *n*=79 | 34.82±9.55 | 15.24±10.70 | 37/42 | 55/24 | 19.58±12.37 |
|  | IGE *n*=0 | — | — | — | — | — |
|  | HC *n*=57 | 34.09±9.69 | — | 30/27 | — | — |
| BRUSSELS | TLE *n*=13 | 35.85±5.27 | 12.50±10.60 | 5/8 | 10/3 | 23.25±13.21 |
|  | IGE *n*=0 | — | — | — | — | — |
|  | HC *n*=44 | 26.64±4.34 | — | 20/24 | — | — |
| CUBRIC | TLE *n*=0 | — | — | — | — | — |
|  | IGE *n*=44 | 27.61±7.39 | 12.95±4.67 | 12/32 | — | 14.61±10.03 |
|  | HC *n*=48 | 28.04±8.17 | — | 14/34 | — | — |
| EPICZ | TLE *n*=40 | 37.73±8.06 | 16.50±13.20 | 18/22 | 17/23 | 21.23±13.00 |
|  | IGE *n*=0 | — | — | — | — | — |
|  | HC *n*=96 | 35.58±8.80 | — | 47/49 | — | — |
| EPIGEN_3.0 | TLE *n*=13 | 40.39±6.28 | 21.84±13.16 | 7/6 | 8/5 | 18.55±11.98 |
|  | IGE *n*=0 | — | — | — | — | — |
|  | HC *n*=64 | 33.11±8.00 | — | 35/29 | — | — |
| GREIFSWALD | TLE *n*=0 | — | — | — | — | — |
|  | IGE *n*=25 | 32.56±7.85 | 19.44±9.73 | 12/13 | — | 13.12±9.96 |
|  | HC *n*=96 | 25.76±5.00 | — | 36/60 | — | — |
| IDIBAPS-HCP | TLE *n*=46 | 35.02±8.43 | 15.22±11.28 | 21/25 | 14/32 | 18.97±10.35 |
|  | IGE *n*=3 | 38.33±3.51 | — | 2/1 | — | — |
|  | HC *n*=51 | 32.75±5.35 | — | 23/28 | — | — |
| KCL_CNS | TLE *n*=13 | 38.85±8.32 | 16.73±13.34 | 4/9 | 5/8 | 22.91±15.80 |
|  | IGE *n*=31 | 29.94±8.42 | 12.00±5.73 | 11/20 | — | 17.94±8.93 |
|  | HC *n*=97 | 30.82±7.39 | — | 45/52 | — | — |
| KUOPIO | TLE *n*=0 | — | — | — | — | — |
|  | IGE *n*=35 | 27.86±8.34 | 17.46±9.93 | 13/22 | — | 10.46±10.67 |
|  | HC *n*=67 | 25.16±1.55 | — | 34/33 | — | — |
| MNI | TLE *n*=79 | 33.10±8.82 | 17.09±10.08 | 34/45 | 44/35 | 16.01±10.72 |
|  | IGE *n*=0 | — | — | — | — | — |
|  | HC *n*=45 | 30.24±6.65 | — | 25/20 | — | — |
| NYU | TLE *n*=17 | 31.76±7.34 | 15.12±7.74 | 7/10 | 7/10 | 17.12±11.68 |
|  | IGE *n*=35 | 32.43±9.34 | 14.23±5.25 | 19/16 | — | 18.38±9.86 |
|  | HC *n*=108 | 28.51±8.88 | — | 52/56 | — | — |
| RMH | TLE *n*=25 | 34.80±9.22 | 23.24±12.72 | 15/10 | 16/9 | 12.03±12.51 |
|  | IGE *n*=21 | 31.33±8.79 | 22.28±9.43 | 8/13 | — | 9.51±13.81 |
|  | HC *n*=19 | 29.21±8.59 | — | 12/7 | — | — |
| UCSD | TLE *n*=17 | 33.24±8.42 | 14.87±12.60 | 8/9 | 10/7 | 19.53±13.53 |
|  | IGE *n*=0 | — | — | — | — | — |
|  | HC *n*=29 | 30.10±7.77 | — | 16/13 | — | — |
| UNAM | TLE *n*=17 | 30.76±9.50 | 14.53±12.73 | 7/10 | 9/8 | 16.18±9.54 |
|  | IGE *n*=0 | — | — | — | — | — |
|  | HC *n*=31 | 31.61±10.58 | — | 7/24 | — | — |
| UNICAMP | TLE *n*=161 | 40.83±7.64 | 11.49±9.70 | 70/91 | 91/70 | 29.65±11.55 |
|  | IGE *n*=37 | 32.27±9.31 | 11.46±7.22 | 9/28 | — | 20.81±10.08 |
|  | HC *n*=357 | 32.26±8.83 | — | 135/222 | — | — |
| UNIMORE | TLE *n*=0 | — | — | — | — | — |
|  | IGE *n*=38 | 24.61±7.46 | 11.32±5.75 | 14/24 | — | 11.39±9.86 |
|  | HC *n*=34 | 28.47±5.25 | — | 14/20 | — | — |
| XMU | TLE *n*=40 | 28.25±8.45 | 17.18±12.06 | 25/15 | 25/15 | 11.28±8.02 |
|  | IGE *n*=11 | 31.00±10.45 | 18.56±16.49 | 7/4 | — | 12.00±10.23 |
|  | HC *n*=13 | 31.54±6.99 | — | 9/4 | — | — |

**TABLE S2.** List of disease-specific risk genes from previously published GWAS.

| Disorder | Genes | Ref |
| --- | --- | --- |
| All epilepsies | ATXN1, BCL11A, BRD7, C3orf33, FANCL, GABRA2, GJA1, GRIK1, HEATR3, KCNAB1, KCNN2, PCDH7, PNPO, SCN1A, SCN2A, SCN3A, STAT4, STX1B, TTC21B, ZEB2 | 32 |
| Focal epilepsy (hippocampal sclerosis) | C3orf33, GJA1, KCNAB1 | 32 |
| Generalized epilepsy | ATXN1, BCL11A, FANCL, GABRA2, GRIK1, KCNN2, PCDH7, PNPO, SCN1A, SCN2A, SCN3A, STAT4, TTC21B | 32 |
| Attention deficit/hyperactivity disorder | ARTN, B4GALT2, CCDC24, DPH2, DUSP6, FOXP2, IPO13, KDM4A, LINC00461, PCDH7, POC1B, PTPRF, SEMA6D, SORCS3, SPAG16, ST3GAL3, TMEM161B_AS1 | 41 |
| Autism spectrum disorder | APOPT1, BAG5, C8orf74, CADPS, CKB, FEZF2, KCNN2, KIZ, KLC1, KMT2E, MACROD2, MROH5, NEGR1, NKX2_2, NUDT12, PINX1, POU3F2, PTBP2, SOX7, TMEM33, TRMT61A, XRN2 | 42 |
| Bipolar disorder | ADCY2, ADD3, ANK3, CACNA1C, CD47, FADS2, FSTL5, GRIN2A, HDAC5, LMAN2L, MRPS33, NCAN, PACS1, PC, PLEKHO1, POU3F2, RIMS1, RPS6KA2, SCN2A, SHANK2, SSBP2, STARD9, STK4, THSD7A, TRANK1, ZCCHC2, ZNF592 | 43 |
| Major depressive disorder | ABT1, ACVR1B, ANKHD1, ANKK1, ANKS1B, AP3B1, APOPT1, ARHGEF25, ASCC3, ASIC2, ASTN2, ASXL3, ATP1A3, BAD, BAG5, BAZ2B, BCHE, BSN, BTN2A1, BTN3A2, BTN3A3, C16orf45, CABP1, CACNA1E, CACNA2D1, CAMKK2, CDH13, CDH22, CDH9, CDK14, CELF2, CELF4, CHD6, CKB, CNTN5, CNTNAP5, CRB1, CSMD1, CTNNA3, CTTNBP2, DCC, DCDC1, DENND1A, DENND1B, DLST, DRD2, ELAVL2, EMILIN3, EPHB2, ERBB4, ESRRG, ETFDH, EXT1, FADS1, FADS2, FAM120A, FAM120AOS, FAM172A, FANCL, FCF1, FH, FHIT, FNIP2, GINM1, GPC5, GPC6, GRIK5, GRM5, GRM8, GTF2IRD1, HARS, HIST1H2BF, HIST1H2BL, HIST1H2BN, HIST1H2BO, HS6ST3, HSPA1A, HTT, IGSF6, IK, INPP4B, KDM3A, KIAA1143, KLC1, KMT2A, KYNU, LIN28B, LPIN3, LRFN5, LRP1B, LST1, LTBP3, MANEA, MAP9, MED19, MEF2C, MEGF11, METTL9, MGAT4C, MICB, MIER1, MR1, MYBPC3, MYRF, NEGR1, NICN1, NRG1, OLFM4, PAX6, PCDH9, PCDHA1, PCDHA5, PCLO, PCNP, PGBD1, PLA2R1, PLCL1, PMFBP1, POGZ, PPID, PPP6C, PRR16, PRSS16, PSEN2, PSORS1C1, PTPRS, RAB27B, RAB3B, RABEPK, RANGAP1, RBFOX1, RBMS3, RFTN2, RHOA, RHOBTB1, RPS6KL1, RSRC1, RTN1, SAMD5, SCAI, SCYL1, SDK1, SEMA6D, SERPING1, SF3B1, SGIP1, SHISA9, SLC4A9, SORBS3, SORCS3, SOX5, SOX6, SPPL3, SPRY2, STAU1, STK19, TAL1, TCAIM, TCF4, TCTEX1D1, TENM2, TMEM106B, TMEM161B, TMEM258, TMEM42, TMEM67, TRAF3, TRMT10C, TRMT61A, TTC12, UBE2M, USP3, VPS41, VRK2, YLPM1, ZC3H7B, ZCCHC7, ZDHHC21, ZDHHC5, ZFHX4, ZHX3, ZKSCAN4, ZKSCAN8, ZMAT2, ZNF165, ZNF184, ZNF322, ZNF35, ZNF536, ZNF638, ZNF660, ZSCAN26, ZSCAN31, ZSCAN9 | 44 |
| Schizophrenia | ABCB9, ACTR5, ADAMTSL3, ALDOA, AMBRA1, ANKRD44,<br>ANP32E, APH1A, APOPT1, ARHGAP1, ARL3, ARL6IP4,<br>ASPHD1, ATG13, ATP2A2, BAG5, C12orf65, C1orf54, C2orf69,<br>CA14, CACNA1C, CACNA1I, CACNB2, CCDC39, CDK2AP1,<br>CENPM, CHADL, CHRM4, CHRNA3, CILP2, CKAP5, CKB,<br>CLCN3, CNKSR2, CNNM2, CNTN4, DGKZ, DNAJC19, DOC2A,<br>DPYD, DRD2, EFHD1, ERCC4, ESAM, FAM57B, FANCL, FES,<br>FUT9, FXR1, GATAD2A, GIGYF2, GNL3, GRM3, HAPLN4,<br>HARBI1, HIRIP3, HSPD1, HSPE1, IMMP2L, INA, IREB2,<br>KCNV1, KCTD13, KDM4A, KLC1, L3MBTL2, LRP1, MAD1L1,<br>MAN2A2, MAPK3, MDK, MPHOSPH9, MSANTD2, MSL2,<br>NAB2, NAGA, NCAN, NCK1, NDUFA13, NDUFA4L2, NDUFA6,<br>NEK1, NEK4, NGEF, NISCH, NRGN, NT5C2, NT5DC2, NXPH4,<br>OGFOD2, OTUD7B, PBRM1, PBX4, PCCB, PCGF6, PITPNM2,<br>PJA1, PLCH2, PLCL1, PLEKHO1, PPP1R13B, PPP1R16B,<br>PPP2R3A, PTPRF, PUS7, R3HDM2, RANGAP1, RFTN2, RILPL2,<br>SBNO1, SEPT3, SEZ6L2, SF3B1, SGSM, SHISA8, SHMT2,<br>SLC32A1, SLC35G2, SLC39A8, SLC45A1, SMDT1, SMG6,<br>SMIM4, SNAP91, SNX19, SREBF2, SRR, STAB1, STAG1, STAT6,<br>SUGP1, TAC3, TAF5, TCF20, TCF4, TM6SF2, TMEM219,<br>TNFRSF13C, TRANK1, TRIM8, TRMT61A, TSNARE1, TSR1,<br>TYW5, VRK2, WBP1L, YPEL3, ZFYVE21, ZSCAN2, ZSWIM6 | 45 |

**FIGURE S1.**
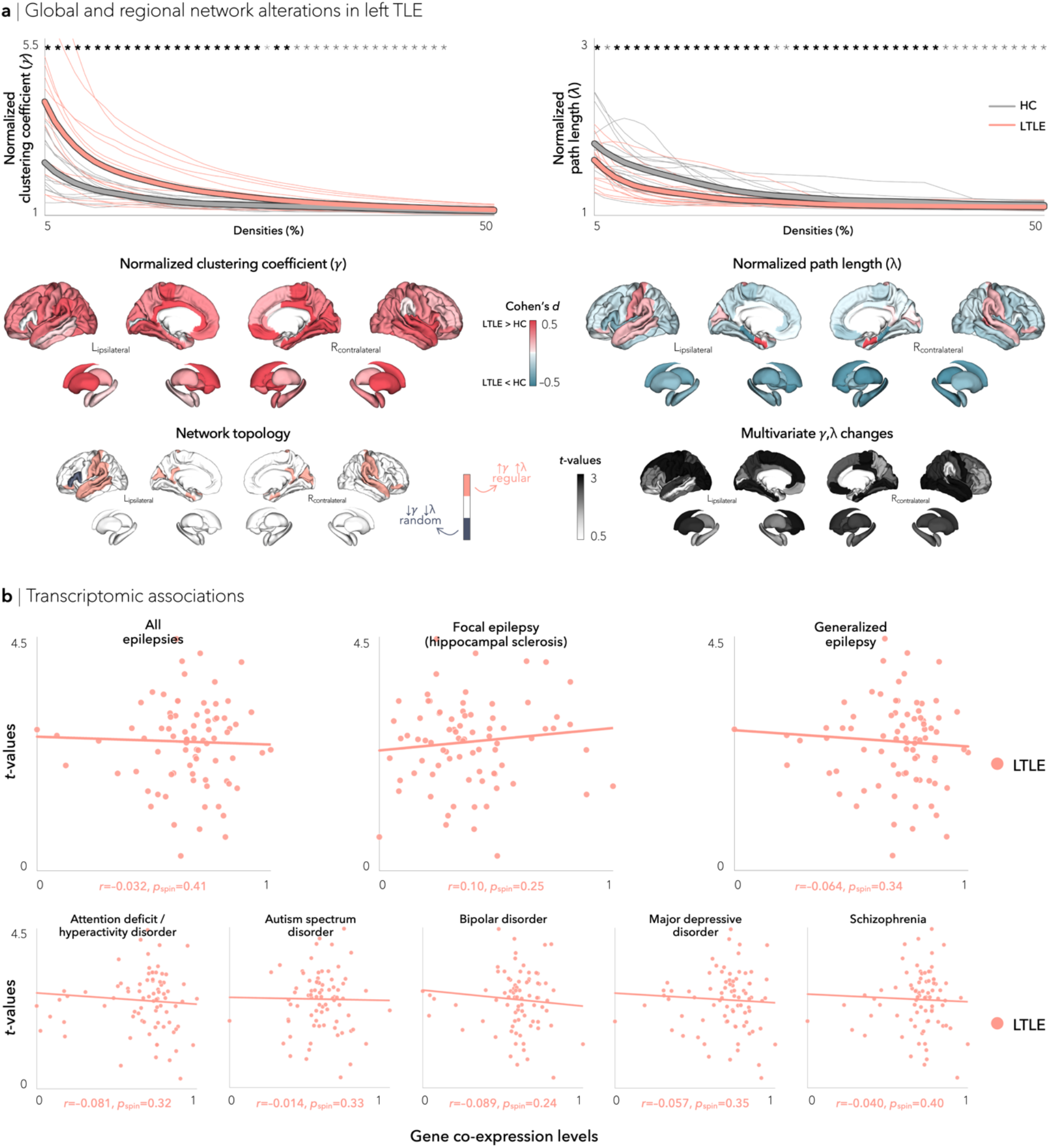
Structural covariance networks in left TLE (LTLE). (**a**) Global differences in clustering coefficient (top left) and path length (top right) between left LTLE and healthy controls (HC) are plotted as a function of network density. Increased small-worldness (increased clustering coefficient, decreased path length) was observed in individuals with left TLE. Student’s t-tests were performed at each density value; bold asterisks indicate *p*_FDR_<0.05, semi-transparent asterisks indicate *p*_FDR_<0.1. Thin lines represent data from individual sites. Multivariate topological differences in left TLE were primarily observed in bilateral fronto-temporal cortices, and revealed a regular network configuration (increased clustering and path length). (**b**) Associations between epilepsy-related gene co-expression profiles and multivariate changes were marginal, although correlation to hippocampal sclerosis genes was strongest.

**FIGURE S2.**
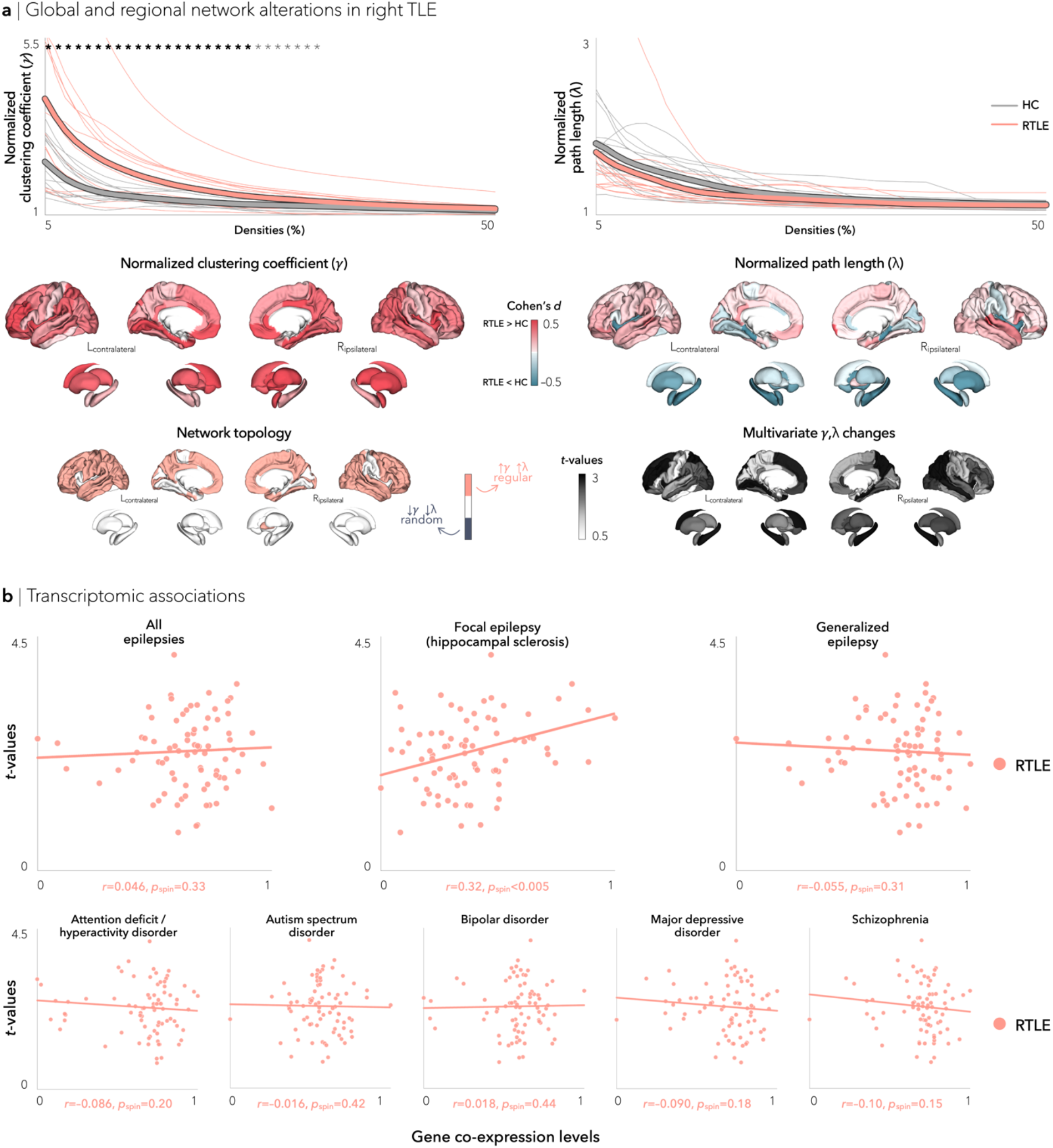
Structural covariance networks in right TLE (RTLE). (**a**) Global differences in clustering coefficient (top left) and path length (top right) between RTLE and healthy controls (HC) are plotted as a function of network density. Increased small-worldness (increased clustering coefficient, decreased path length) was observed in individuals with right TLE. Student’s t-tests were performed at each density value; bold asterisks indicate *p*_FDR_<0.05, semi-transparent asterisks indicate *p*_FDR_<0.1. Thin lines represent data from individual sites. Multivariate topological changes in right TLE were primarily observed in bilateral fronto-temporal cortices and the hippocampus, and revealed a widespread regular network configuration (increased clustering and path length). (**b**) Associations between epilepsy-related gene co-expression profiles and multivariate changes were strongest for hippocampal sclerosis genes.

**FIGURE S3.**
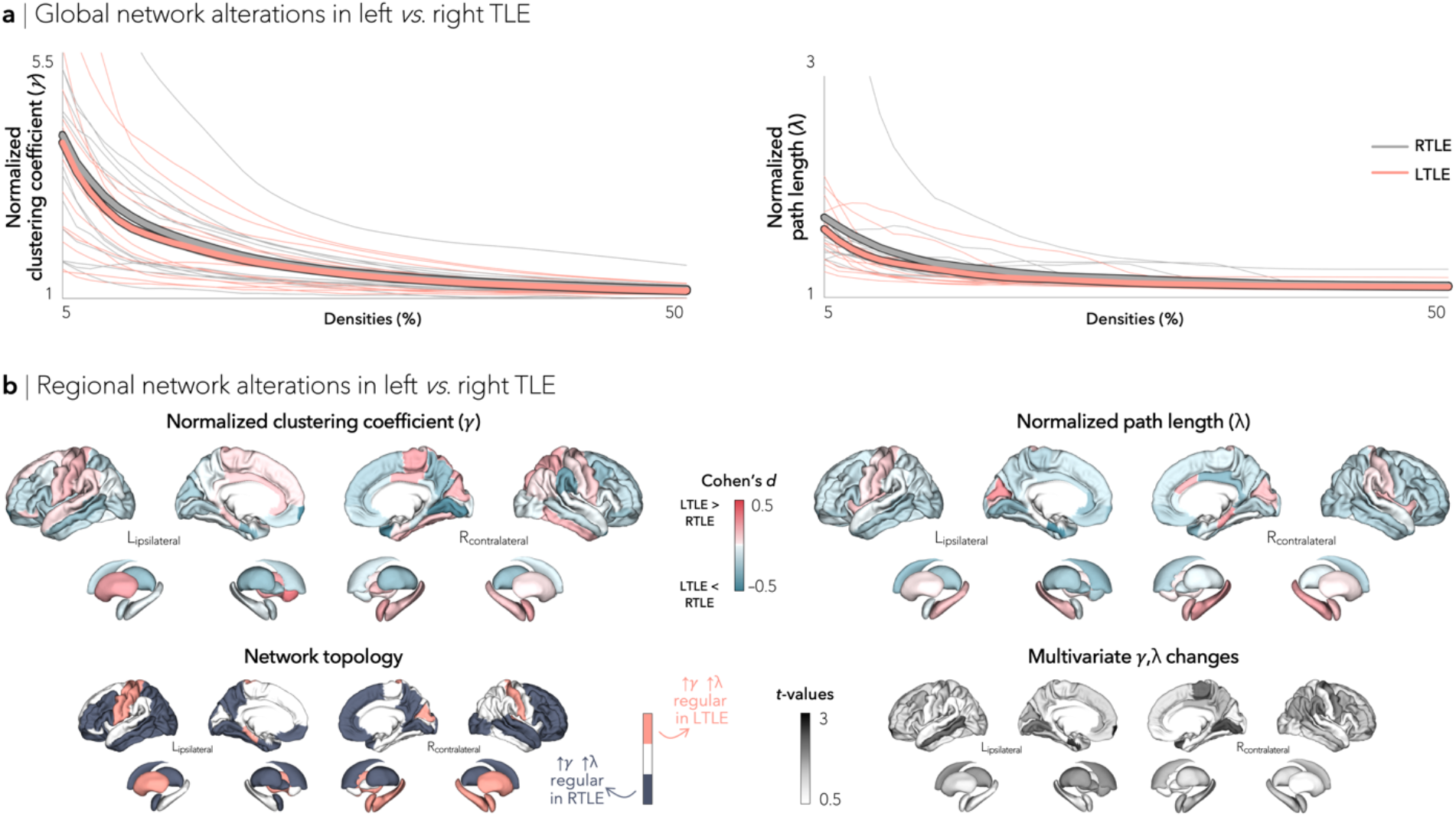
Structural covariance networks in left (LTLE) *vs*. right (RTLE) TLE. (**a**) Global differences in clustering coefficient (left) and path length (right) between LTLE and RTLE are plotted as a function of network density. No significant difference was observed. Student’s t-tests were performed at each density value; bold asterisks indicate *p*_FDR_<0.05, semi-transparent asterisks indicate *p*_FDR_<0.1. Thin lines represent data from individual sites. (**b**) Trends for multivariate topological changes in LTLE *vs*. RTLE were observed in ipsilateral middle frontal gyrus and enthorinal cortex as well as contralateral calcarine sulcus. Compared to the other subcohort, LTLE showed network regularization (increased clustering and path length) in sensorimotor cortices, whereas RTLE showed widespread network regularization.

**FIGURE S4.**
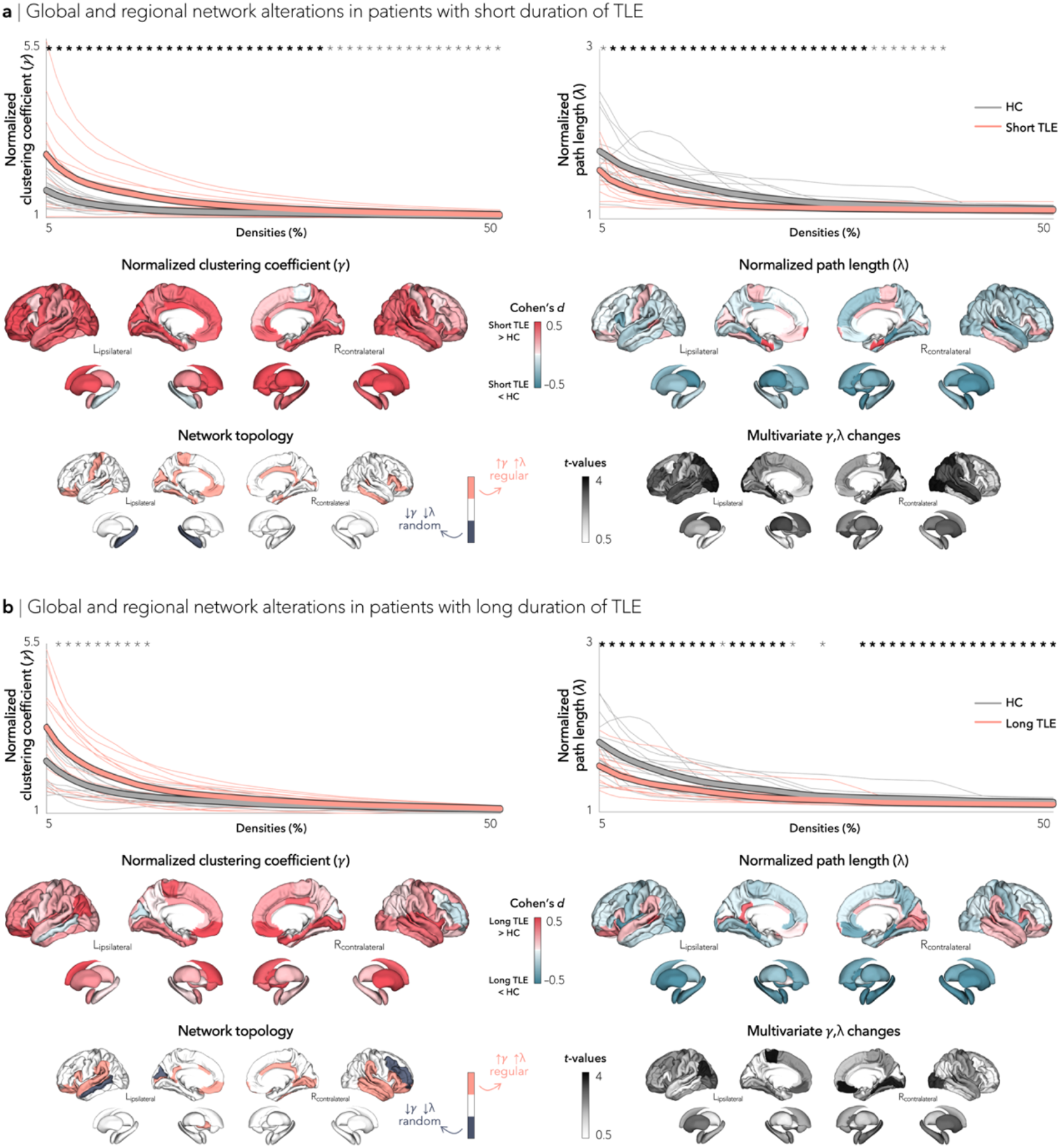
Structural covariance networks in patients with short and long TLE duration. (**a**) Global differences in clustering coefficient (top left) and path length (top right) between Short TLE and healthy controls (HC) are plotted as a function of network density. Increased small-worldness (increased clustering coefficient, decreased path length) was observed in patients with short TLE duration. Student’s t-tests were performed at each density value; bold asterisks indicate *p*_FDR_<0.05, semi-transparent asterisks indicate *p*_FDR_<0.1. Thin lines represent data from individual sites. Multivariate topological changes in patients with short TLE duration were widespread, affecting bilateral fronto-parietal and limbic cortices and the thalamus, and revealed a regular subnetwork configuration (increased clustering and path length). (**b**) Global differences in clustering coefficient (top left) and path length (top right) between Long TLE and healthy controls (HC) are plotted as a function of network density. Increased small-worldness (increased clustering coefficient, decreased path length) was observed in patients with long TLE duration. Multivariate topological changes in patients with long TLE duration were primarily observed in fronto-limbic cortices and bilateral putamen, amygdala, and hippocampus, and revealed a regular subnetwork configuration (increased clustering and path length).

**FIGURE S5.**
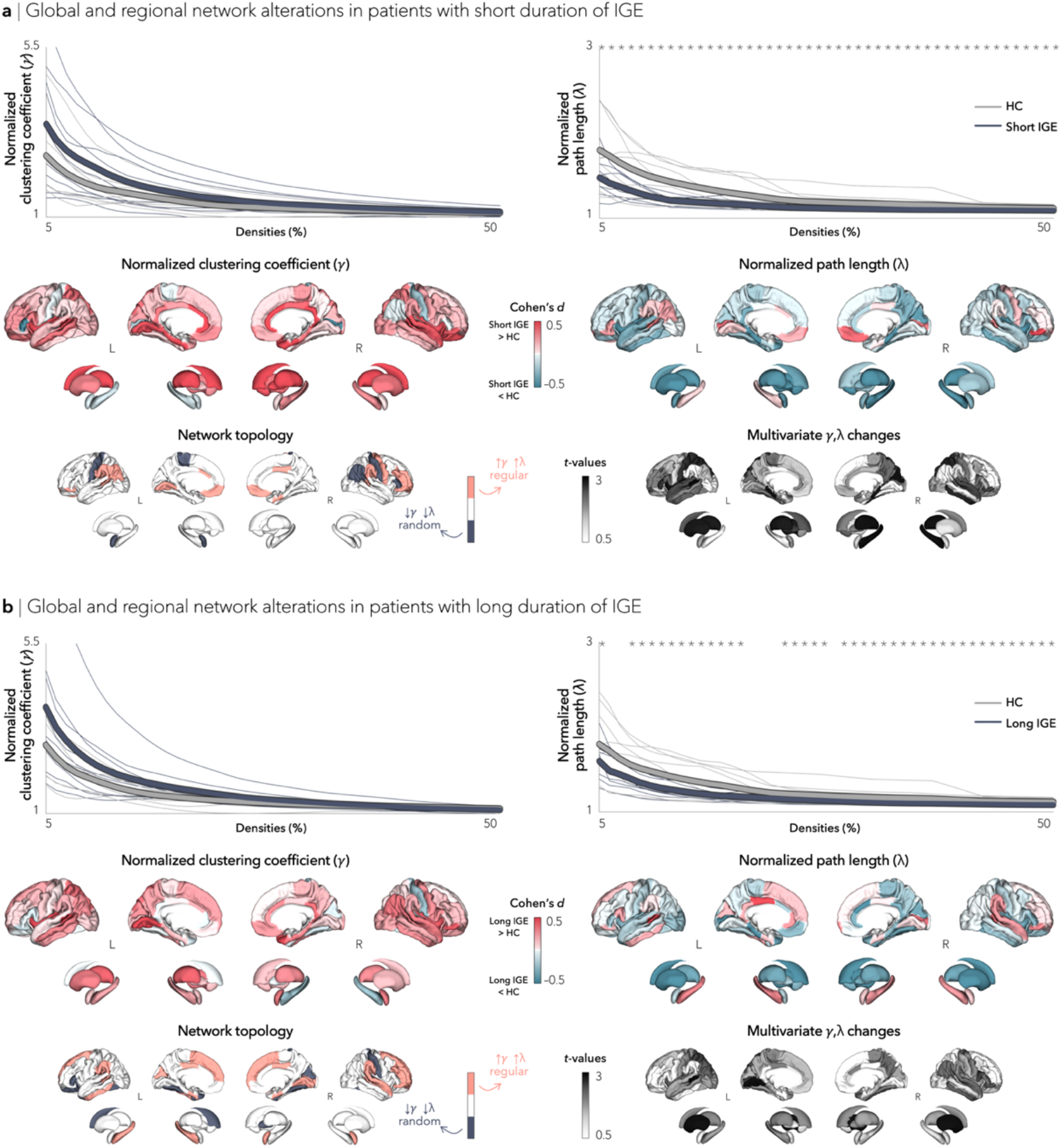
Structural covariance networks in patients with short and long IGE duration. (**a**) Global differences in clustering coefficient (top left) and path length (top right) between Short IGE and healthy controls (HC) are plotted as a function of network density. Decreased path length was observed in patients with short IGE duration. Student’s t-tests were performed at each density value; bold asterisks indicate *p*_FDR_<0.05, semi-transparent asterisks indicate *p*_FDR_<0.1. Thin lines represent data from individual sites. Multivariate topological changes in patients with short IGE duration were observed, primarily in bilateral fronto-central and temporal cortices, as well as bilateral thalamus and right hippocampus. Both random (decreased clustering and path length in centro-parietal regions) and regular (increased clustering and path length in orbitofrontal and parietal regions) network configurations were observed. (**b**) Global differences in clustering coefficient (top left) and path length (top right) between Long IGE and healthy controls (HC) are plotted as a function of network density. Decreased path length was observed in patients with long IGE duration. Multivariate topological changes in patients with long IGE duration were primarily observed in bilateral fronto-temporo-parietal regions and bilateral putamen. Once again, both random (fronto-central cortices) and regular (fronto-temporo-parietal regions) network configurations were observed.

**FIGURE S6.**
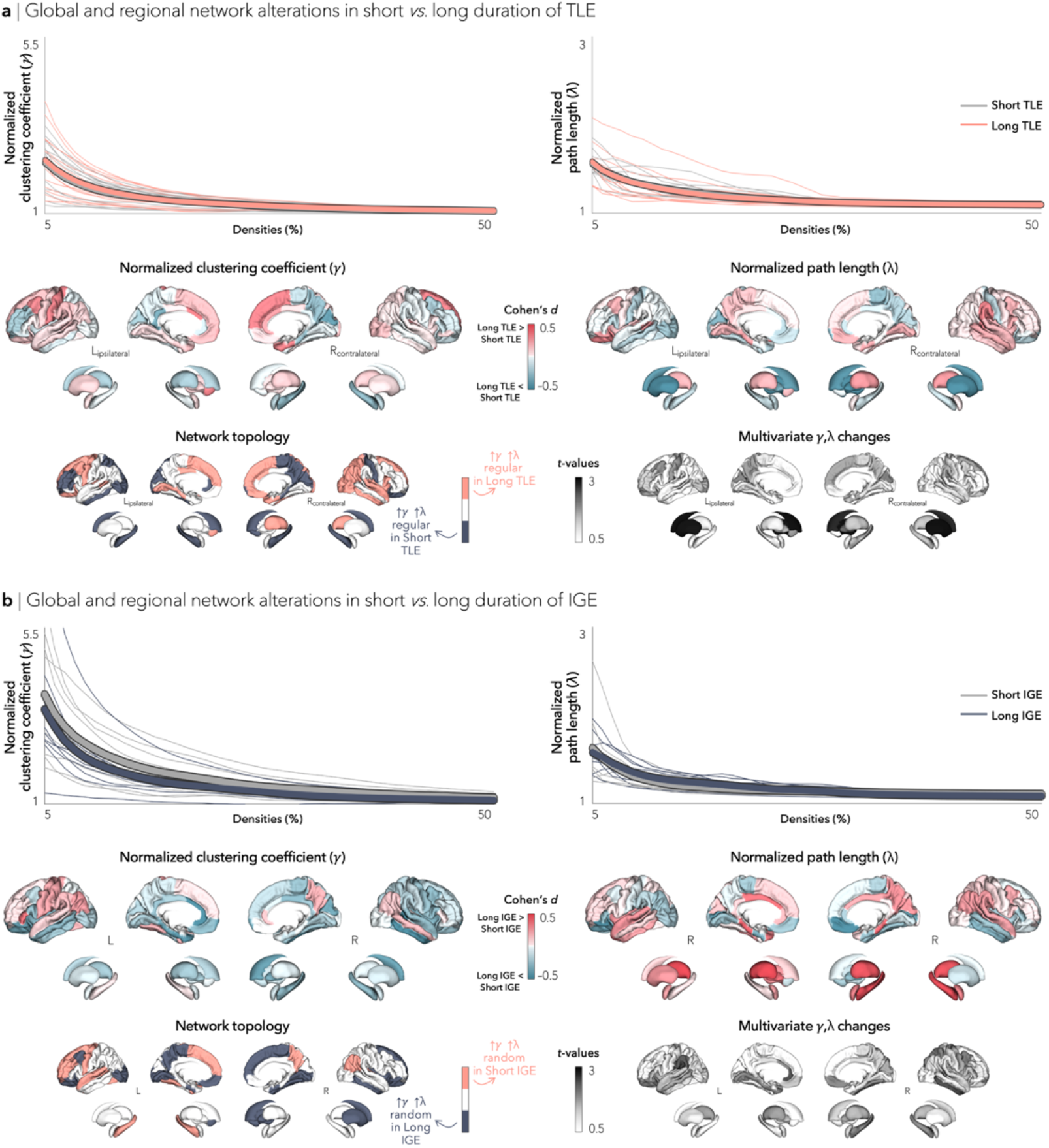
Structural covariance networks in short *vs*. long TLE/IGE duration. (**a**) Global differences in clustering coefficient (top left) and path length (top right) between Short and Long TLE are plotted as a function of network density; no significant difference was observed in clustering coefficient or path length. Student’s t-tests were performed at each density value; bold asterisks indicate *p*_FDR_<0.05, semi-transparent asterisks indicate *p*_FDR_<0.1. Thin lines represent data from individual sites. Marginal multivariate topological changes were observed between groups, primarily affecting bilateral caudate and ipsilateral putamen. Patients with shorter duration exhibited network regularization (increased clustering and path length) in bilateral fronto-parietal cortices and hippocampus, whereas patients with longer duration exhibited network regularization more broadly in fronto-temporo-parietal cortices. (**b**) Global differences in clustering coefficient (top left) and path length (top right) between Short and Long IGE are plotted as a function of network density; no significant difference was observed in clustering coefficient or path length. Marginal multivariate topological changes were observed between groups, primarily affecting bilateral fronto-limbic cortices. Patients with shorter duration exhibited network randomization (decreased clustering and path length) in left fronto-temporal cortices and hippocampus and right parietal cortices, whereas patients with longer duration exhibited network randomization in left fronto-parieto-occipital cortices and right fronto-temporal cortices, caudate, and putamen.

**FIGURE S7.**
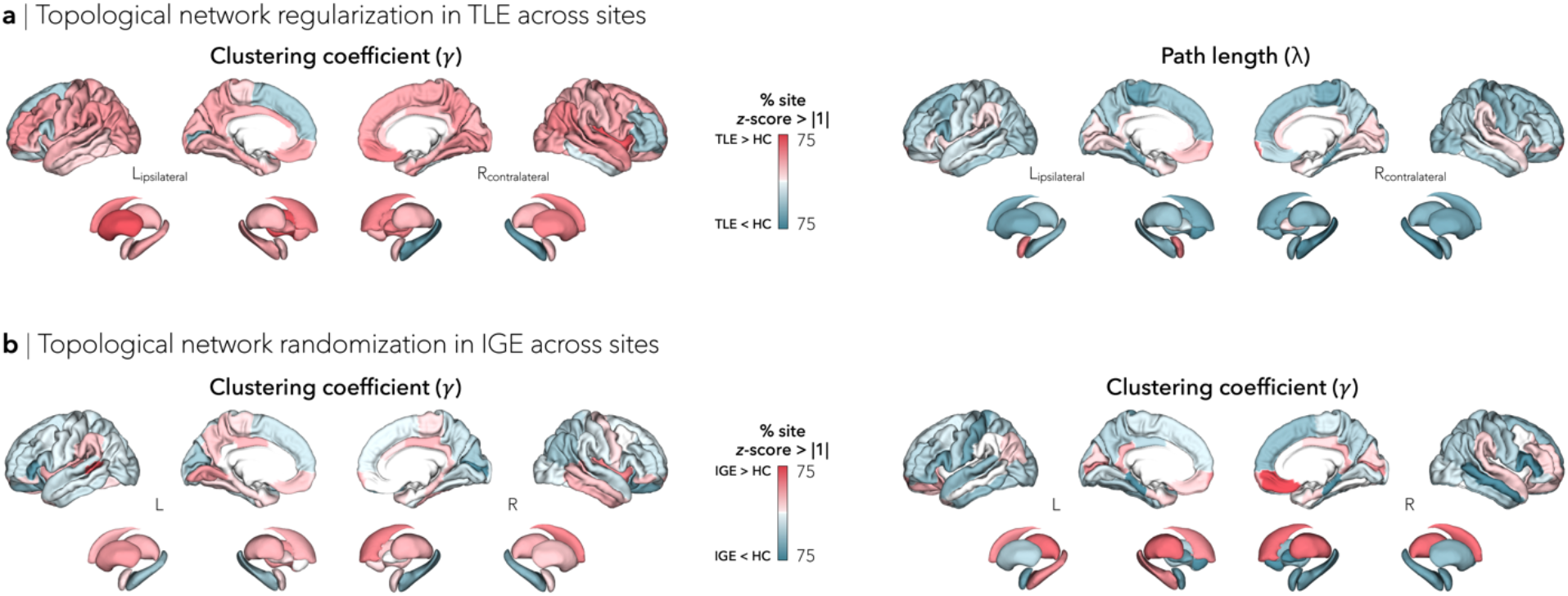
Site-specific structural covariance changes in TLE and IGE. Structural covariance and graph theoretical analyses were repeated in each site independently and yielded virtually identical results. (**a**) In TLE, increased clustering and path length were observed in bilateral orbitofrontal, temporal, and angular cortices, as well as ipsilateral amygdala, revealing a regularized, “lattice-like,” subnetwork. (**b**) In IGE, widespread decreased in clustering and path length were observed in fronto-temporo-parietal regions, suggesting a randomized network configuration.

**FIGURE S8.**
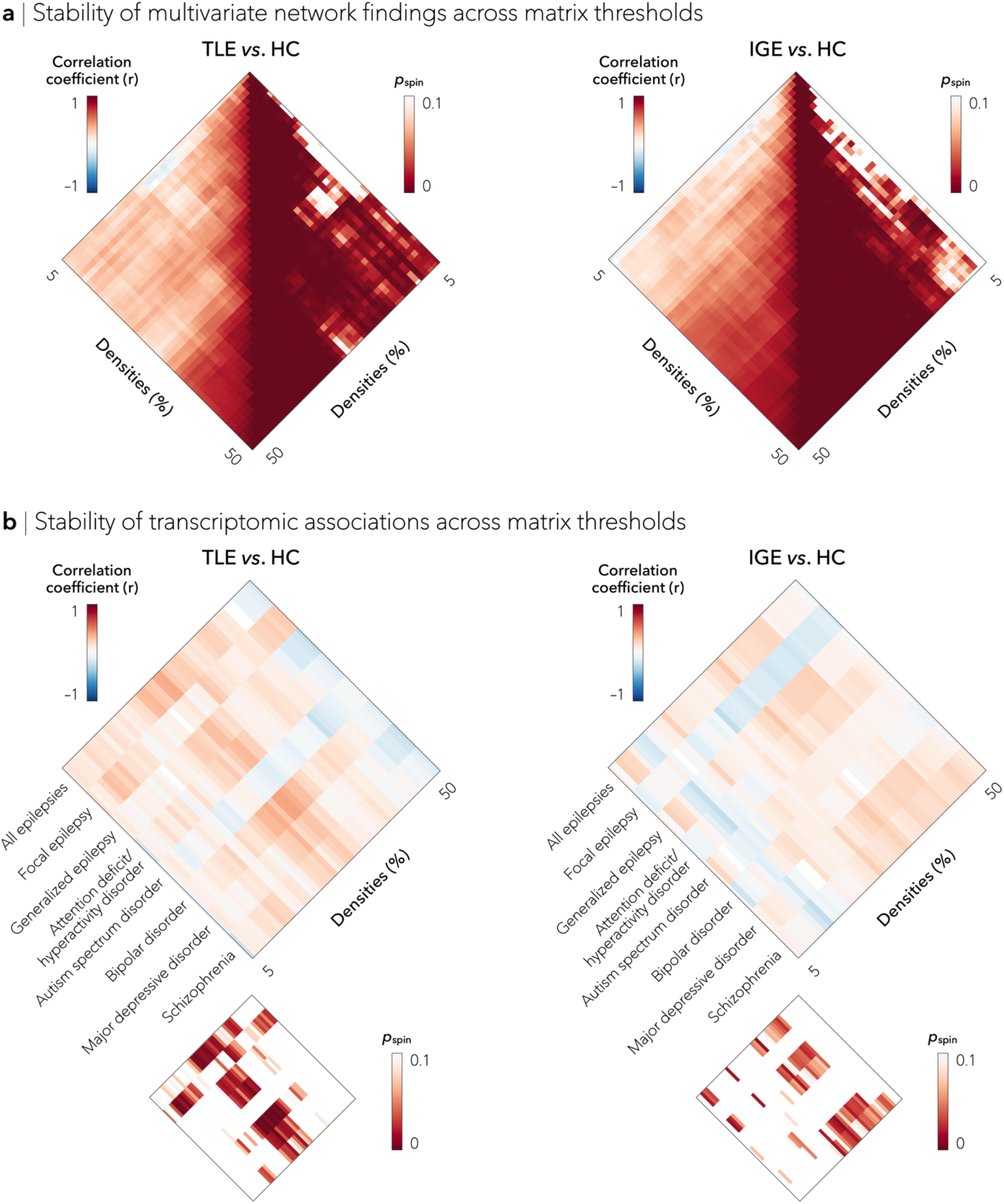
Reproducibility of findings across matrix thresholds. (**a**) Associations between multivariate topological changes (clustering and path length) across matrix thresholds. (**b**) Associations between disease-related gene co-expression maps and multivariate findings across matrix thresholds.

